# Contact Lens Wear Alters Transcriptional Responses to *Pseudomonas aeruginosa* in Both the Corneal Epithelium and the Bacteria

**DOI:** 10.1101/2024.12.03.626720

**Authors:** Naren G. Kumar, Melinda R Grosser, Stephanie Wan, Daniel Schator, Eugene Ahn, Eric Jedel, Vincent Nieto, David J. Evans, Suzanne M. J. Fleiszig

**Affiliations:** Herbert Wertheim School of Optometry & Vision Science, University of California, Berkeley, CA USA; Graduate Program in Infectious Diseases and Immunity, University of California, Berkeley, CA USA; College of Pharmacy, Touro University California, Vallejo, CA USA; Graduate Groups in Vision Science and Microbiology, University of California, Berkeley, CA USA

## Abstract

**Purpose:** Healthy corneas resist colonization by virtually all microbes yet contact lens wear can predispose the cornea to sight-threatening infection with *Pseudomonas aeruginosa*. Here, we explored how lens wear changes corneal epithelium transcriptional responses to *P. aeruginosa* and its impact on bacterial gene expression.

**Methods:** Male and female C57BL/6J mice were fitted with a contact lens on one eye for 24 h. After lens removal, corneas were immediately challenged for 4 h with *P. aeruginosa*. A separate group of naïve mice were similarly challenged with bacteria. Bacteria-challenged eyes were compared to uninoculated naive controls as was lens wear alone. Total RNA-sequencing determined corneal epithelium and bacterial gene expression.

**Results:** Prior lens wear profoundly altered the corneal response to *P. aeruginosa*, including: upregulated pattern-recognition receptors (*tlr3, nod1*), downregulated lectin pathway of complement activation (*masp1*), amplified upregulation of *tcf7*, *gpr55, ifi205, wfdc2* (immune defense) and further suppression of *efemp1* (corneal stromal integrity). Without lens wear, *P. aeruginosa* upregulated mitochondrial and ubiquinone metabolism genes. Lens wear alone upregulated *axl, grn, tcf7, gpr55* (immune defense) and downregulated Ca2^+^-dependent genes *necab1, snx31 and npr3*. *P. aeruginosa* exposure to prior lens wearing vs. naïve corneas upregulated bacterial genes of virulence (*popD*), its regulation (*rsmY*, PA1226) and antimicrobial resistance (*arnB*, *oprR*).

**Conclusion:** Prior lens wear impacts corneal epithelium gene expression altering its responses to *P. aeruginosa* and how *P. aeruginosa* responds to it favoring virulence, survival and adaptation. Impacted genes and associated networks provide avenues for research to better understand infection pathogenesis.

## Introduction

While contact lens wear is a common form of vision correction that is generally well tolerated, it can cause serious complications such as corneal infection (1), most commonly caused by *Pseudomonas aeruginosa* (2,3). Such infections can be severe, and can cause permanent vision impairment (1,3,4).

When healthy, the corneal epithelium resists bacterial adhesion even if challenged with large inocula of *P. aeruginosa* or other pathogens (5,6). This intrinsic resistance to bacterial adhesion has been studied by us and others (7), our own work showing that it requires MyD88 and associated surface receptors IL- 1R and TLR4 (6,8–10), resident CD11c+ cells (9) and TRPA1 and TRPV1 ion channels associated with corneal sensory nerves (11). Known mediators of this resistance include antimicrobial peptides (12–16), surfactant proteins (17,18) and membrane-associated and secreted mucins (19–21). Some of these factors are found in tear fluid, a mucosal fluid that despite containing factors that have antimicrobial activity fails to directly kill many strains of *P. aeruginosa* (22). However, tear fluid does play important roles in defense, acting in tandem with other intrinsic defenses against this pathogen to bolster epithelial defenses and altering gene expression in the bacteria (18,22–29).

Less is known about how these defenses are compromised by lens wear (7). Obstacles to progress include ethical limitations around performing infection research in people, and technical/practical limitations surrounding animal models of lens wear. With respect to the latter, contact lens wear can predispose animals to corneal infection, including rabbits, rats and mice (30–33) showing their potential. While an advantage of using rabbits is that they can be fitted with human lenses, this generally involves suturing their eyelids closed to retain the lens. Their size, expense and the limited availability of reagents for rabbits has also limited their utility. Similar problems exist for rat models, which do not fit human lenses. While a plethora of reagents are available for mouse research, manufacturing lenses to fit them is challenging due to the small eye size and shape of their corneas. Thus, our current understanding of contact lens infection pathogenesis is largely derived from cell culture experiments, use of animal infection models without lens wear, and correlative/observational/epidemiological studies of lens wearing people, which have been used creatively by many scientists. Factors thought to be involved in lens related infection pathogenesis include a disruption to tear exchange or tear function (34,35), compromise to epithelial barrier function (36), reduced epithelial proliferation (37), suppression of antimicrobial peptide expression (38), bacterial biofilm formation on lenses and bacterial adaptations on posterior lens surfaces (30) and trapping of host immune cells and associated factors (31). Our own studies support hypotheses for how these factors conspire to compromise defense. For example, we have shown that outer membrane vesicles (OMV’s) are released by *P. aeruginosa* in response to prolonged tear fluid exposure (e.g. under a lens) and that these can kill epithelial cells on the surface of mouse eyes, then enabling susceptibility to *P. aeruginosa* adhesion (39). In another study, we showed that corneal epithelial cells shed from human subjects were more susceptible to *P. aeruginosa* adhesion after contact lens wear (40). This method was later used by others to show a role for hypoxia in this increased bacterial adhesion (41). Those findings supported approval of silicone hydrogel lens materials with high oxygen permeability which unfortunately did not reduce the incidence of infection (42), a finding further supported by later lens-wear studies in a rabbit model (31), thereby questioning the role of hypoxia in human lens-related infections.

More recently, we developed a mouse model for contact lens wear that does not require lid suturing to retain the lens on the eye and demonstrated that it mimics multiple features of human lens wear (32). For example, we showed that they are comfortable for mice to wear, do not cause detectable loss of corneal epithelial barrier function to fluorescein, and during wear the become colonized with the same type of Gram-positive bacteria as lenses worn by humans (32). Also reported to be a feature of human lens wear, they induced a parainflammatory (sub-clinical) response involving changes to immune cell numbers, morphology and location (32), which can persist for several days after lens removal (43). Importantly, they too predispose to infection with *P. aeruginosa*, the resulting pathology showing features similar to human lens related infections. This model enables use of the full range of modern research tools available for use in mice, allowing for detailed mechanistic studies not currently possible using humans or other species.

Here, we used bulk RNA-sequencing to explore how prior lens wear in mice changes the corneal epithelial response to *P. aeruginosa* challenge compared to how it responds when it is naïve to lens wear. The experimental design additionally provided insights into two other related questions; how prior lens wear alone (without bacteria) impacts gene expression in the cornea, and how prior lens wear changes the bacterial response to the cornea.

## Materials and Methods

### Bacteria

*Pseudomonas aeruginosa* strain PAO1, originally sourced from the University of Washington (44) was grown on tryptic soy agar plates at 37 °C for ∼16 h. Inocula were prepared by suspending bacteria in PBS to a concentration of ∼10^11^ CFU/ml, as confirmed by viable counts.

### Mouse Lens Wear Model and Experimental Groups

All procedures involving animals were carried out in accordance with the standards established by the Association for the Research in Vision and Ophthalmology, under a protocol (AUP-2019-06-12322-1) approved by the Animal Care and Use Committee, University of California Berkeley. This protocol adheres to PHS policy on the humane care and use of laboratory animals, and the guide for the care and use of laboratory animals.

Male and female six-week-old C57BL/6J mice were fitted with a custom-made silicone hydrogel contact lens on one eye as previously described (32). Prior to lens fitting, mice were anesthetized using 1.5 - 2% isoflurane delivered via precision vaporizer (VetEquip Inc., Pleasanton, CA) A Handi-Vac suction pen (Edmund Optics, Barrington, NJ) with a 3/32″ probe was used for contact lens handling and fitting as previously explained in detail (30,32). Lens wearing mice were fitted with an Elizabethan collar (Kent Scientific) and single-housed without enrichments to prevent lens removal. Pure-o’Cel paper bedding (The Andersons Inc., Maumee OH) was used to reduce dust levels in the cage. Mice were allowed to wear the contact lenses for 24 h, after which they were checked for lens retention using a stereomicroscope (Zeiss, Stemi 2000-C) while under brief isoflurane anesthesia. Mice that lost their contact lens were excluded from further experimentation. Non-lens wearing control mice were handled similarly and also wore an Elizabethan collar over the same time period.

Fig. 1 shows the experimental set-up with 4 groups each containing 3-4 mice. Group 1) No lens wear, no bacterial inoculation; Group 2) Lens wear for 24 h, no bacterial inoculation; Group 3) No lens wear, then bacterial inoculation for 4 h; Group 4) Lens wear for 24 h, lens removed then bacterial inoculation for 4 h. Since the primary goal of the study was to determine how prior lens wear impacts the cornea’s response to bacterial inoculation, with both the comparison and control groups inoculated (the variable being prior lens wear status), sham inoculation was not needed. Prior to bacterial inoculation, mice were anesthetized with ketamine (80 - 100 mg/Kg) and dexmedetomidine (0.25 - 0.5 mg/Kg), lenses were removed if applicable, and corneas inoculated with 5 µl of a ∼10^11^cfu/ml bacterial suspension, re- inoculating every hour for a total incubation period of 4 h (4 inoculations in total). During exposure to bacteria, mice remained anesthetized and covered under a heat lamp for the entire 4 h period. Mice were then euthanized with a lethal dose of ketamine-xylazine (80-100 mg/Kg and 5-10mg/Kg respectively) followed by cervical dislocation. The experiment was repeated once.

**Figure 1:**
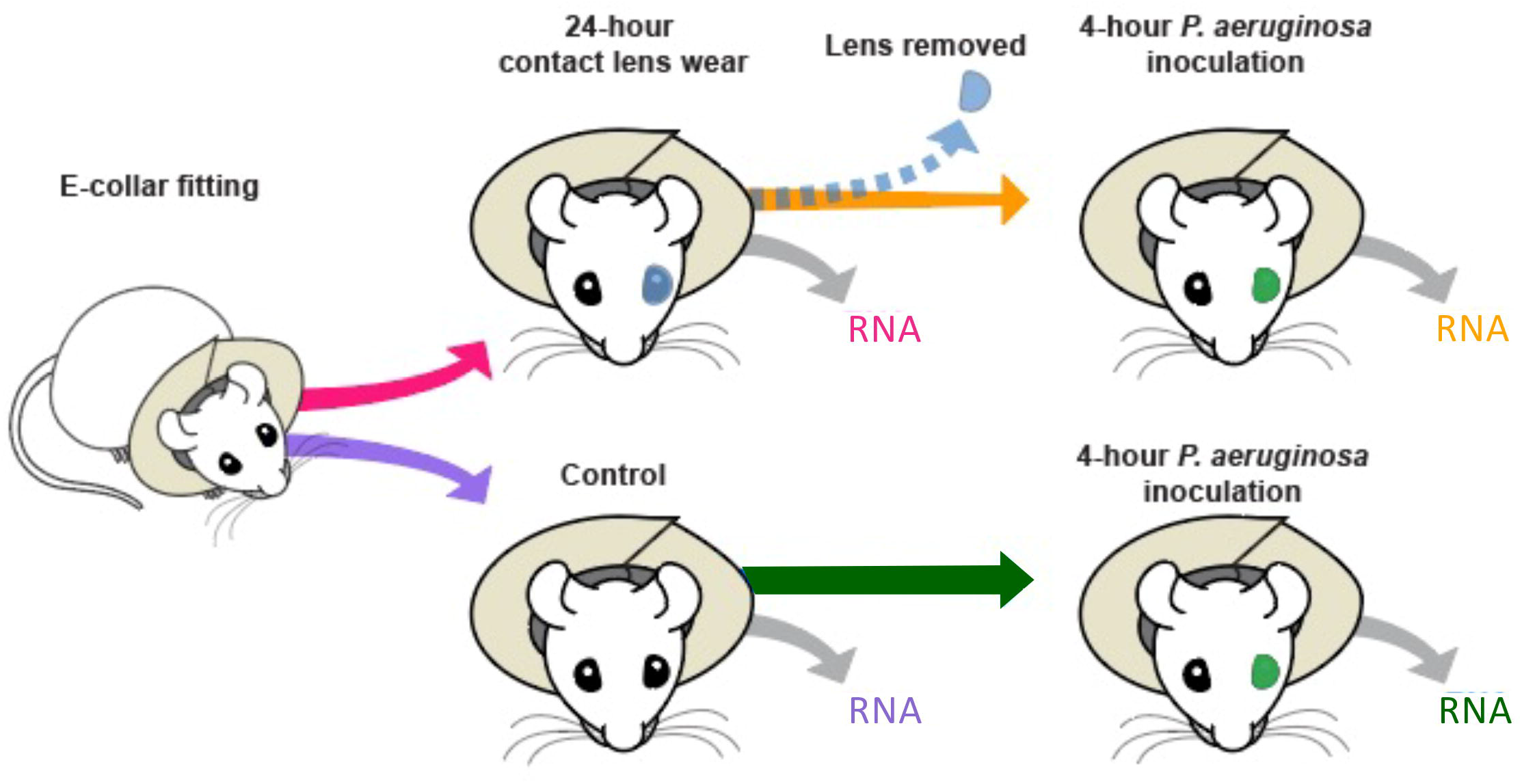
Schematic diagram of experimental design. The lens wear group of mice was separate from the naïve control group. *P. aeruginosa* (green) was inoculated after lens wear or to naïve controls in a 5 µl drop of a 10^11^cfu/ml suspension. Over a 4 h exposure period, bacterial inoculation was performed 4 times.

### RNA Extraction

Immediately after euthanasia, the corneal surface was rinsed three times with PBS and the eye excised. The corneal epithelium was then collected for RNA extraction using an Alger brush with a 0.5 mm burr (Gulden Ophthalmics, Elkins Park, PA) while observing under a stereomicroscope to ensure precise collection. The collected epithelium was removed from the Alger brush by moistening in sterile PBS, then placement into 1 ml of ice-cold Tri-Reagent and Tough Micro-Organism Lysing Mix with 0.5 mm glass beads (Omni International, Kennesaw GA). Cells from all mice in each group were pooled after collection for the tissue disruption step using an Omni Bead-Ruptor. The pooled and disrupted tissue was then stored at -80° C until RNA extraction. RNA was extracted directly from the TRI reagent using a Direct-Zol RNA Mini-Prep Kit according to manufacturer instructions (Zymo Research Corporation, Irvine, CA). RNA from all four groups was processed using a Ribo-Zero rRNA Removal Kit (Human/Mouse/Rat). RNA from the two groups that had been inoculated with bacteria were additionally processed with a Ribo-Zero rRNA Removal Kit (Bacteria) (Illumina, San Diego, CA). A Kapa Biosystems library preparation kit was used to prepare a standard-sized library with custom Unique Dual indexes. Libraries were sequenced on a NovaSeq 6000 platform with 50 bp single reads. Raw reads will be deposited on the SRA (Sequence Read Archive) NCBI (https://www.ncbi.nlm.nih.gov/sra) upon acceptance for publication (Accession #).

## Data Processing

Raw reads were mapped to their respective genomes as previously described (45). Host reads were aligned to mouse genome (*Mus_musculus*.GRCm39.109.gtf) and bacterial reads to the *Pseudomonas aeruginosa* genome (*Pseudomonas_aeruginosa_pao1_gca_000006765.ASM676v1.56.gtf*). Reads were trimmed with Trimmomatic. FastQ Screen and MultiQC were used to determine the percentage of mapped reads and aggregate results. For mouse reads, the genome index for each genome was built using HISAT2 and sorted and indexed using SamTools. HTSeq was used to create a read count matrix and annotate genes. After identifying raw reads from the host genome, unmapped reads were mapped onto the bacterial genome using Bowtie2 and a count matrix was created using HTSeq. CombatSeq was used to detect and correct batch effects. Low quality read counts were filtered from all samples (raw count filter > 10). DESeq2 was used to determine differentially expressed genes with a full model with all samples. Differentially- expressed genes were determined using the general formula with modifications: *[results (full, cooksCutoff = FALSE, independentFiltering = TRUE, lfcThreshold=.5, altHypothesis="greaterAbs", contrast = c("Group","PAO1_4h","Control"))]*.

### Analysis of Differentially-Expressed Genes and Networks

Differentially-expressed genes were determined using DESeq2 workflow (46). Network analysis was performed using Cytoscape and STRING Enrichment was used to obtain a reduced list of biologically relevant genes for each condition. All differentially-expressed genes with P < 0.01 and log_2_Fold-Change (FC) > 0.5 were selected and imported into Cytoscape. GENEMANIA was used to obtain the base gene network and converted to a STRING network using ENSEMBL gene ID. STRING Enrichment was performed with a term redundancy cut-off of 0.5. Enrichment Map (*Jaccard similarity* > 0.4) was used on the set of enriched terms to plot the enriched categories with a unique identifier FDR < 0.05. The network was organized using Attribute Grid Layout using *Log2FoldChange* as the criteria. Possible transcriptional factors or genes whose change due to a treatment has the greatest impact on other genes were identified using *ClosenessCentrality* cutoff > 0.4.

### Enrichment of Immune-Related Genes

ClueGo was used to identify immune pathway related genes differentially-expressed in the dataset. Differentially-expressed genes P < 0.01 and Log_2_FC > 0.5 were selected and imported into Cytoscape. ClueGo was performed on ENSEMBL gene ID and Network specificity score of 0.75 and using “GO Term fusion”. Batch-corrected values of genes associated with immune pathways were filtered and plotted using ggplot2. ClueGo clusters were detected using Global analysis and number of genes per cluster = 5.

## Results

### Corneal Epithelium RNA-Sequencing Analysis

MutliQC plots were utilized to show the percentage of reads per sample that mapped to the mouse genome (Table 1). The read percentage of single mapping reads to the mouse genome ranged between 14.77% to 59.43% and only single-mapping reads were used for downstream analysis. Differential gene expression analysis was performed separately for corneal epithelium and bacteria, the latter addressed separately below. Unsupervised analysis of host gene expression profiles using Principal Component Analysis (PCA) showed that each group could be distinguished from the others and principal components PC1 and PC2 contributed to 40% and 33% variance respectively (Fig. 2). Gene expression profiles of mice wearing contact lenses for 24 h (Fig. 2, Magenta) were clearly distinguished from naive controls (Fig. 2, purple). Exposure to *P. aeruginosa* for 4 h post 24 h contact lens wear (Fig. 2, orange) or for 4 h to naïve corneas (Fig. 2, green) further distinguished the gene expression profiles from uninoculated corneas with prior lens wear impacting the corneal epithelial response to bacterial challenge.

**Figure 2.**
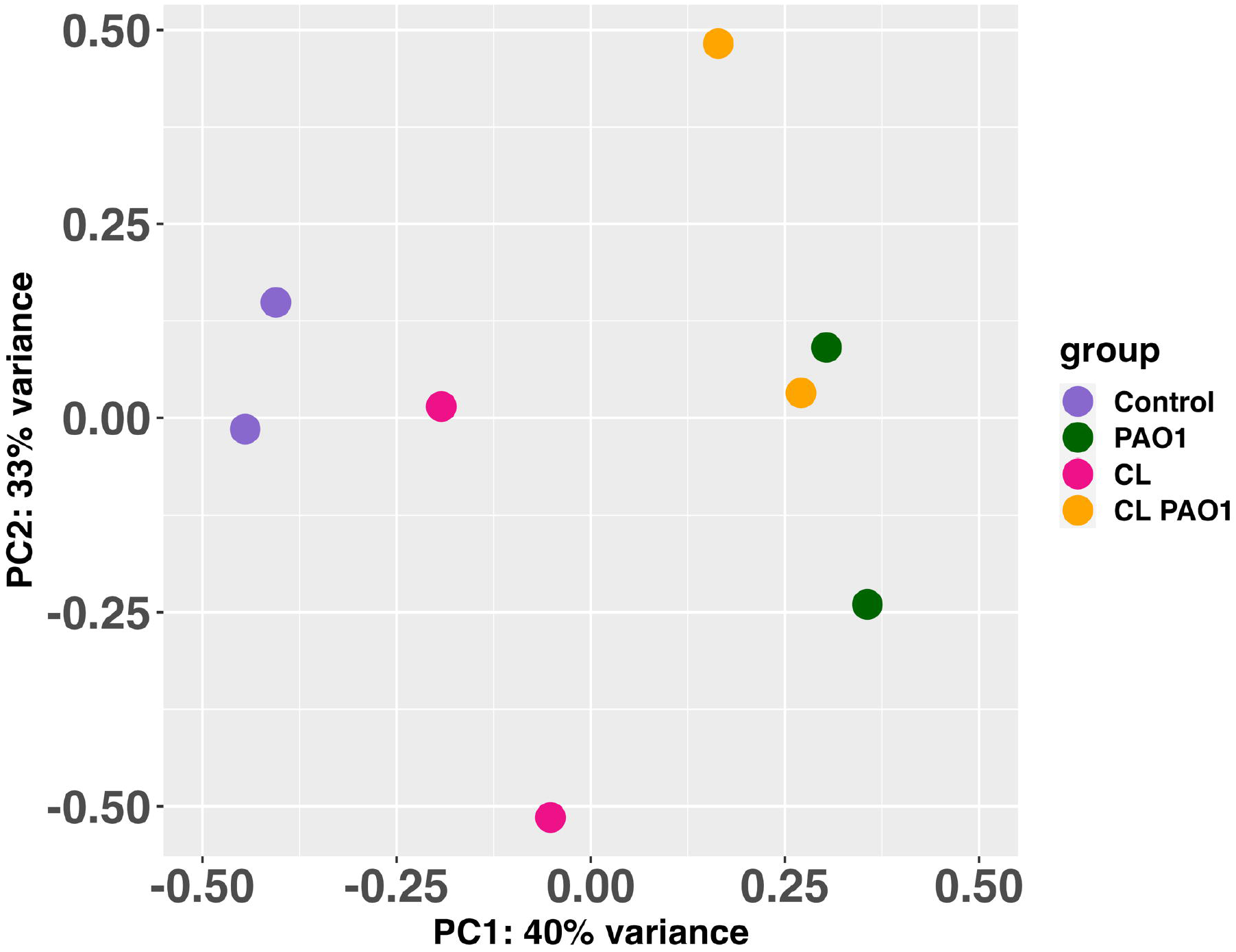
PCA analysis showing distinct gene expression profiles for each RNA-sequencing sample.

**Table 1:** RNA sequencing host read alignment statistics to the mouse genome (GRCm39)

| **#Fastq_screen**<br>**Version: 0.15.3** | **#Aligner:**<br>**Bowtie2** | **#Reads in**<br>**Subset:**<br>**100000** |  |  |  |  |
| --- | --- | --- | --- | --- | --- | --- |
| **Sample ID** | **Group** | **#**<br>**Reads**<br>**Processed** | **#**<br>**Unmapped** | **%**<br>**Unmapped** | **#**<br>**One Hit**<br>**One**<br>**Genome** | **%**<br>**One Hit**<br>**One**<br>**Genome** |
| 1-Collar_only_1_S147_<br>L006_R1_001_screen | Control | 100102 | 35496 | 35.46 | 49266 | 49.22 |
| 2-Collar_only_2_S148_<br>L006_R1_001_screen | Control | 99938 | 35718 | 35.74 | 47839 | 47.87 |
| 3-CL_1_S149_<br>L006_R1_001_screen | CL | 99998 | 30151 | 30.16 | 31183 | 31.18 |
| 4-CL_2_S150_<br>L006_R1_001_screen | CL | 99960 | 23503 | 23.52 | 59406 | 59.43 |
| 5-PAO1_1_S151_<br>L006_R1_001_screen | PAO1 | 99967 | 52461 | 52.48 | 22880 | 22.89 |
| 6-PAO1_2_S152_<br>L006_R1_001_screen | PAO1 | 100023 | 25353 | 25.34 | 59149 | 59.14 |
| 7-CL PAO1_1_S153_<br>L006_R1_001_screen | CL-PAO1 | 100007 | 66614 | 66.61 | 14775 | 14.77 |
| 8-CL PAO1_2_S154_<br>L006_R1_001_screen | CL-PAO1 | 100075 | 51797 | 51.75 | 21272 | 21.26 |

### Impact of Prior Lens Wear on the Corneal Transcriptome Response to *P. aeruginosa*

We asked how prior contact lens wear impacted the corneal epithelial transcriptomic response to *P. aeruginosa*. The rationale was to gain insights into how contact lens wear predisposes the corneal epithelium to infection susceptibility. To this end, we compared the profile of differential gene expression with and without prior lens wear when corneas were exposed to *P. aeruginosa*, in both cases versus baseline naïve corneas not exposed to a lens or bacteria. Fig. 3A shows a Venn diagram of differential gene expression after bacterial exposure for the lens wear and non-lens wear groups (Groups 4 and 3, respectively) each versus naïve corneas that had not worn a lens or been inoculated (Group 1). A total of 498 genes were deregulated by *P. aeruginosa* challenge compared to completely naïve corneas. Of these, 143 were deregulated irrespective of whether the cornea had worn a lens. Another 224 were deregulated only if a lens had been worn, the majority of them upregulated (189 genes) rather than downregulated (35 genes) (Log_2_FC > 1, P < 0.01). This differed from the 131 genes impacted in only the no previous lens wear group, with 55 upregulated and 76 downregulated (Log_2_FC > 1, P < 0.01).

**Figure 3.**
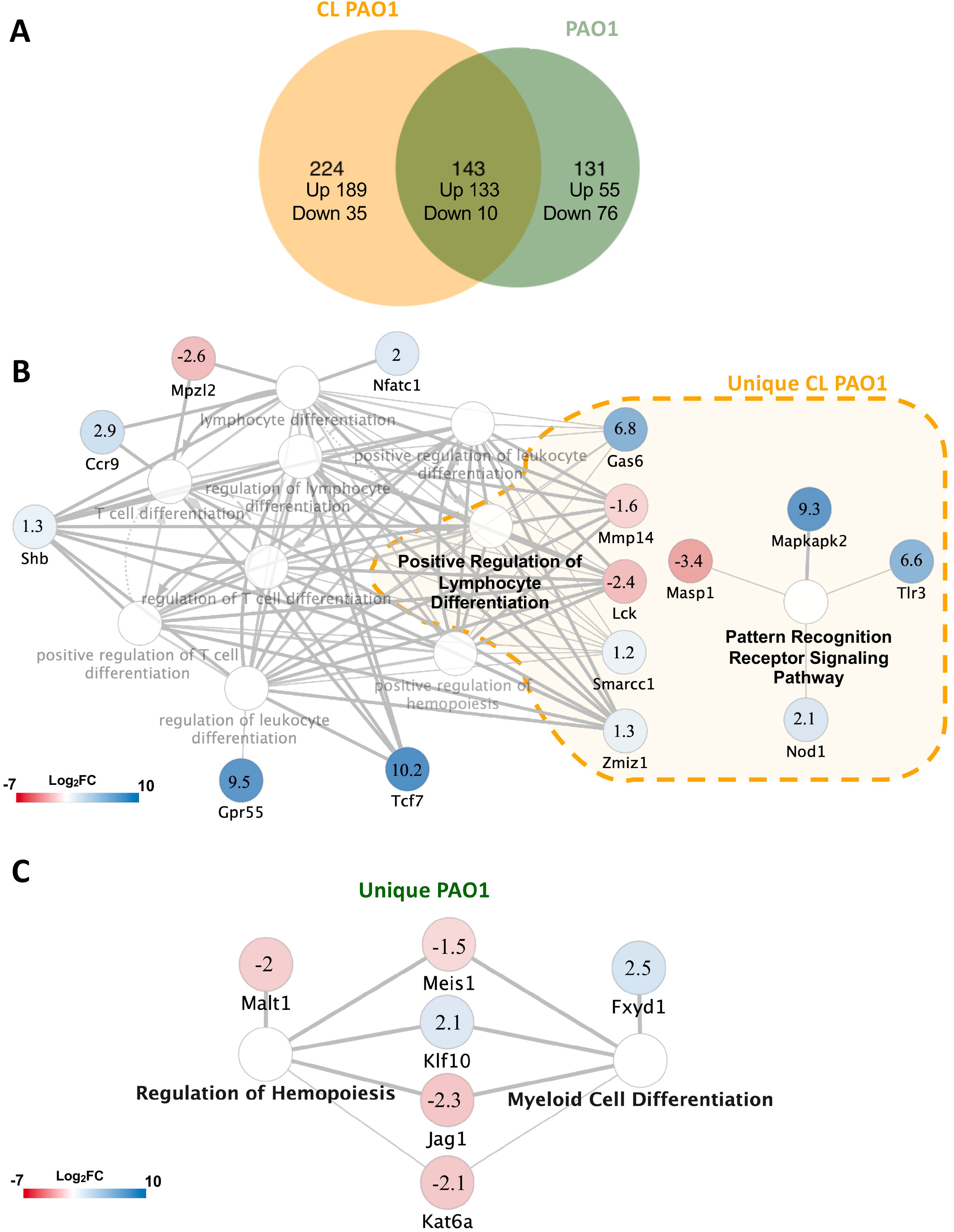
Mouse cornea epithelium transcriptomic response to *P. aeruginosa* challenge. A) Venn diagram showing differentially-expressed genes after 4 h *P. aeruginosa* challenge to a prior contact lens wearing cornea (CL PAO1) or to a naive cornea (PAO1 only). Numbers indicate deregulated genes unique to CL- PAO1 (orange) or to PAO1 (green) or common to both conditions (overlap). B) ClueGO immune pathway network for the 367 deregulated genes in CL PAO1 relative to the naive control (no bacteria). The dashed orange box denotes a network of genes unique to CL PAO1 (224 total, complete list in Supplemental Table S1). Deregulated genes outside the dashed orange box overlap with the comparison of PAO1 only versus naïve cornea (143 total, complete list in Supplemental Table S2A). Nodes are annotated with Log_2_ Fold-Change CL PAO1 relative to naive control. C) ClueGO analysis of 131 deregulated genes unique to a comparison of PAO1 only versus naïve control (no bacteria). Nodes annotated with the Log_2_ Fold- Change for that comparison: Blue = upregulated, Red = downregulated in their respective comparisons.

Fig. 3B shows the ClueGO network analysis of the 367 differentially-expressed genes in the cornea after lens wear followed by *P. aeruginosa* challenge versus naïve controls exposed to neither, which highlighted involvement of immune regulatory networks. The analysis identified 15 deregulated genes associated with four major clusters: “Pattern recognition receptor signaling pathway”, “Positive regulation of lymphocyte differentiation”, “Positive regulation of hemopoiesis” and “Regulation of leukocyte differentiation”, highlighting the combined impact of prior contact lens wear (24 h) and *P. aeruginosa* challenge (4 h) on immune cell pathway gene expression. Sub-group analysis using only the 224 deregulated genes that were altered by *P. aeruginosa* challenge only if a lens had been worn (i.e. not if there was no prior lens wear) identified 9 deregulated genes within two clusters: “Pattern recognition receptor signaling pathway” and “Positive regulation of lymphocyte differentiation”. Notable changes within these clusters included, upregulation of *tlr3* (Log_2_FC 6.6), *nod1* (Log_2_FC 2.1) and *mapkapk2* (Log_2_FC 9.3) all involved in immune recognition of antigens including those associated with recognizing bacteria, and downregulation of *masp1* (Log_2_FC -3.4), which is involved in activation of the complement system *via* the lectin pathway (47,48), involved in defense against microbes including *P. aeruginosa*. Also of note was the upregulation of *gas6* (Log_2_FC 6.8), an anti-inflammatory regulator of TLR signaling (49), and down-regulation of *lck*, a lymphocyte-specific tyrosine kinase, which plays a key role in T cell receptor signaling, T cell activation and homeostasis, and other lymphocyte functions (50). A complete list of 224 deregulated genes comparing CL PAO1 with naïve cornea is shown in Supplemental Table S1.

Fig. 3B also shows an analysis of the 143 genes that were deregulated by *P. aeruginosa* irrespective of whether the cornea had worn a lens. While no ClueGO immune networks were detected in the sub- analysis of these common genes, the combined analysis revealed 6 distinct deregulated genes of interest (see Fig. 3B, outside dashed orange box). They included upregulation of *tcf7* (Log_2_FC 10.2), the T cell- specific transcription factor, *gpr55* (Log_2_FC 9.5) involved in neutrophil recruitment in response to injury (51) and *ccr9* encoding a chemokine receptor expressed on numerous immune cells with both proinflammatory and immunoregulatory functions (52,53). It also revealed the down-regulation of *mpzl2* (Log_2_FC -2.6), an epithelial junctional protein. The complete list of 143 deregulated genes within the comparison of CL PAO1 with naïve cornea that overlapped with the comparison of *P. aeruginosa* with naïve cornea can be found in Supplemental Table S2A. For these 143 genes deregulated in response to *P. aeruginosa* irrespective of whether a lens had been worn, we examined if there were magnitude differences in the response as a result of prior lens wear. Table 2 shows the relative impact of lens wear on the response of the corneal epithelium to 4 h *P. aeruginosa* challenge, i.e. comparing epithelium response to *P. aeruginosa* in a naïve versus the prior lens wear (24 h) cornea. There were 36 deregulated genes with some notable changes labeled with an asterisk in Table 2: 1) Prior lens wear was associated with greater upregulation of *tcf7*, *gpr55*, *wfdc2* and *ifi205* in response to *P. aeruginosa* than occurred in the naïve cornea, i.e. prior lens wear amplified the corneal epithelium response to bacteria (relative fold-changes of 15.23, 4.13, 3.99 and 2.83 respectively). Each of those 4 genes are known to be associated with innate immune responses to pathogens or their antigens. 2) Prior lens wear further suppressed *efemp1* expression in response to *P. aeruginosa* (relative fold-change of 0.17). The *efemp1* gene encodes Fibulin-3, an extracellular matrix glycoprotein associated with tumor suppression (54) that is highly expressed in the cornea and required for stromal integrity (55). Supplemental Table S2B shows additional information (including P values) regarding these 36 genes differentially-deregulated by *P. aeruginosa* challenge depending on whether a lens had been worn.

**Table 2.**
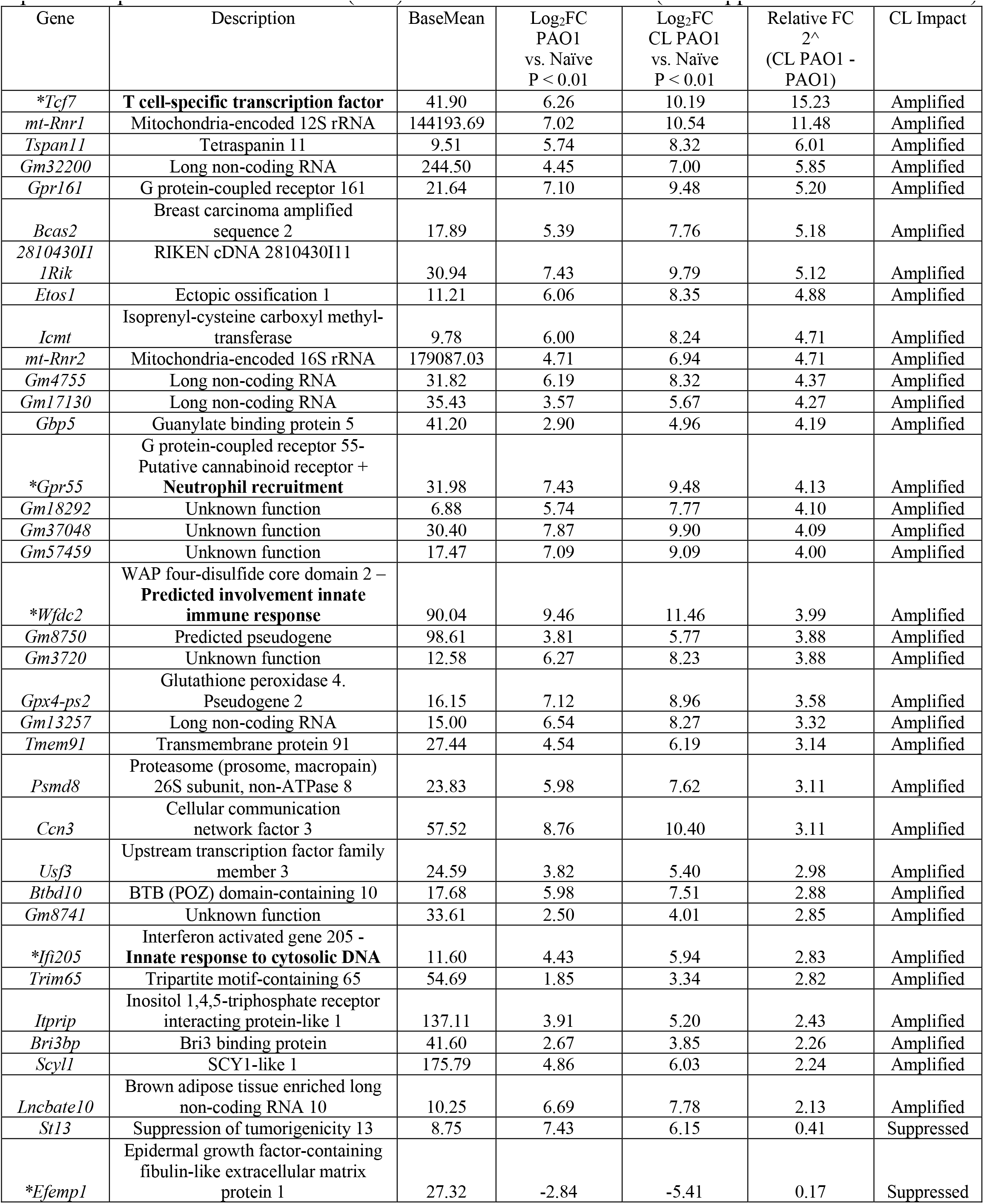
Differential expression of corneal epithelium genes 4 h after *P. aeruginosa* challenge comparing responses of prior lens wear corneas (24 h) to those of naive corneas (see Supplemental Table S2A+B)

Fig. 3C shows a separate analysis of immune pathways involved in the response to bacteria alone using the 131 genes differentially-expressed after *P. aeruginosa* challenge in only the no prior lens wear group (55 upregulated, 76 downregulated, see Fig. 3A). This identified 6 genes associated with two major clusters: “Myeloid Cell Differentiation”, and “Regulation of Hemopoiesis”. The full list of these differentially-expressed genes is shown in Supplemental Table S3. Notable changes included: upregulation of the transcription factor *klf10* (Log_2_FC 2.1) with multiple functions including regulation of cell differentiation (56), and of *fxyd1*(Log_2_FC 2.5) which protects vasculature against oxidative stress (57). Several genes were down-regulated by *P. aeruginosa* exposure including, a lysine acetyl-transferase *kat6A* (Log_2_FC 2.1) (58) and *jag1*(Log_2_FC 2.3), a ligand for Notch signaling and required for normal lens development in the eye (59).

Network analysis approaches such as “closeness centrality” identified putative transcription factors that may mediate corneal epithelium responses to *P. aeruginosa*. Known transcription factors were detected, e.g*. tcf7*, *rasgrf1* (Supplemental Table S2). Top categories from enrichment analysis are shown in Supplemental Fig. S1A and included, ubiquinone metabolism (KW-0830), mitochondrial chromosome (GOCC:0000262) and Parkinson’s Disease (mmu05012). As the ubiquinone metabolism functional group was the most significant category (P < 1.2e^-4^), a closer look at genes in this category was performed. Supplemental Fig. S1B shows a network map of identified genes that included, upregulated mitochondrial genes [*mt-nd1* (Log_2_FC 1.61)*, mt-nd2* (Log_2_FC 2.18)*, mt-nd4* (log_2_FC 1.61) and *mt-nd5* (Log_2_FC 1.91)], and downregulated genes [*irf2bp-1* (Log_2_FC -1.91)*, zfp518a* (Log_2_FC - 1.8)*, esrra* (Log_2_FC -1.83) and *hyls1* (Log_2_FC -1.51)]. These genes were identified as nearest neighbors of genes involved in the ubiquinone network. ClueGo analysis was used to determine if any of these differentially-expressed genes were associated with inflammatory or immune responses. No direct immune regulatory clusters were found. However, changes in genes associated with neurotrophin binding and protein stabilization networks were identified and included the transcription factor *hap1* (Supplemental Fig. S2).

### Contact Lens Wear Alone Impacts the Corneal Epithelium Transcriptome

We next examined the impact of lens wear alone on the corneal epithelial transcriptome. Fig. 4 shows that lens wear had a significant effect on epithelial gene expression with multiple deregulated genes and networks detected. In response to lens wear, 94 genes were differentially-expressed, 90 were upregulated and 4 downregulated. The full list of gene is provided in Supplemental Table S4. Top-differentially- expressed genes shown in Fig. 4A include upregulation of *gpr55* (Log_2_FC 8.6) (mentioned above) and involved in neutrophil recruitment in response to injury, *axl* (Log_2_FC 2.3) encoding a TAM receptor protein tyrosine kinase important for tissue homeostasis and suppression of inflammation (60), and *grn* encoding progranulin, a regulator of lysosome function (Log_2_FC 6.79). Several calcium-dependent genes were downregulated, *necab1* (Log_2_FC -1.33), *snx31* (Log_2_FC -1.05) and *npr3* (Log_2_FC; -1.3). Fig. 4B shows an enrichment map network of differentially-expressed genes involved in the epithelium response to 24 h of lens wear (gene sets are shown in Supplemental Table S5). Fig. 4C shows ClueGo analysis of immune response-related genes involved in the corneal epithelium response to 24 h lens wear alone. A cluster associated with “Regulation of Leukocyte Differentiation” was identified [upregulated *axl*, *grp55*, *klf10*, *tcf7* and downregulated *inpp5d*] (See also Supplemental Table S6).

**Figure 4.**
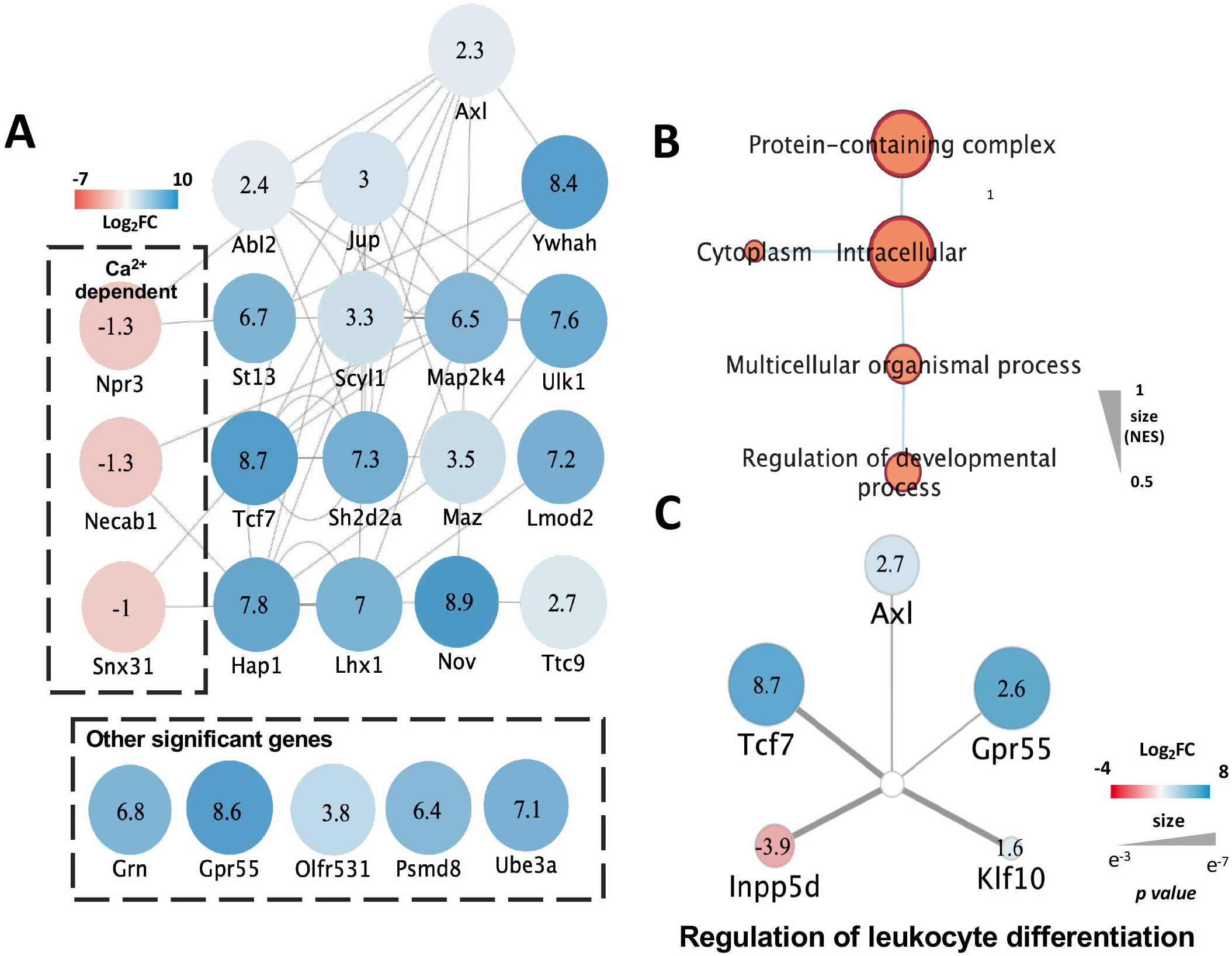
Transcriptional changes in the corneal epithelium after 24 h contact lens wear relative to naive control. A) Top differentially-expressed genes and putative transcription factors identified by network analysis (see Methods and Supplemental Table S4). Nodes are annotated with the Log_2_ Fold-Change. Blue = upregulated, Red = downregulated. B) Enrichment map network of differentially-expressed genes involved in the corneal epithelium response to 24 h lens wear. Node size is proportional to enrichment score (NES= 0.5 – 1) (gene sets listed in Supplemental Table S5). C) ClueGo analysis immune response- related genes. Nodes are annotated with the Log_2_ Fold-Change. Blue = upregulated, Red = downregulated (see Supplemental Table S6).

### Gene expression in bacteria is also differentially-impacted dependent on whether the cornea had previously worn a lens

MutliQC plots were utilized to represent the percentage of reads per sample that mapped to the genome of *P. aeruginosa* strain PAO1 (*Pseudomonas_aeruginosa_pao1_gca_000006765.ASM676v1.56.gtf*). As expected, bacterial reads mapping to the *P. aeruginosa* genome were detected only on eyes challenged with bacteria (Table 3). A total of 183 bacterial genes were differentially-expressed between exposure to corneas that had worn a lens and those that had not. The complete list is shown in Supplemental Table S7, and the 42 most differentially-expressed genes are shown in Table 4 (22 upregulated, 20 downregulated). Notable upregulated genes based upon prior known functions in *P. aeruginosa* pathogenesis include PA1226 (Log_2_FC 5.16) encoding a putative transcriptional regulator and *rsmY* (Log_2_FC 1.55) encoding a small regulatory RNA known to be a global regulator of virulence (61,62). Others include *popD* (Log_2_FC 5.30) a type three secretion system (T3SS) translocon pore protein (63–66), the T3SS being among the most important virulence mechanisms of *P. aeruginosa* (66–70), and *arnB* Log_2_FC 5.14) part of the operon encoding resistance to polymyxin B and other cationic antimicrobial peptides (71–73) and *oprR* (Log_2_FC 6.18) associated with resistance to quaternary ammonium compounds (74) that are commonly used as disinfectants and preservatives in contact lens and other ophthalmic solutions. Bacterial genes most down- regulated after exposure to prior lens wearing corneas encoded “hypothetical proteins” of as yet unknown function, including PA1975 (Log_2_FC -3.79) and PA1168 (Log_2_FC -2.45): others included the *flp* gene encoding type IVb pili (Log_2_FC -1.21).

**Table 3.** Bacteria read alignment statistics *P. aeruginosa* genome (ASM676v1)

| #Fastq_screen<br>Version: 0.15.3 | #Aligner:<br>Bowtie2 | # Reads<br>in Subset:<br>100000 |  |  |  |  |
| --- | --- | --- | --- | --- | --- | --- |
| Sample ID | Group | #<br>Reads<br>Processed | #<br>Unmapped | %<br>Unmapped | #<br>One Hit<br>One<br>Genome | %<br>One Hit<br>One<br>Genome |
| 1-Collar_only_1_S147_<br>L006_R1_001_screen | Control | 100102 | 100088 | 99.99 | 0 | 0 |
| 2-Collar_only_2_S148_<br>L006_R1_001_screen | Control | 99938 | 99927 | 99.99 | 0 | 0 |
| 3-CL_1_S149_<br>L006_R1_001_screen | CL | 99998 | 99951 | 99.95 | 0 | 0 |
| 4-CL_2_S150_<br>L006_R1_001_screen | CL | 99960 | 99945 | 99.99 | 0 | 0 |
| 5-PAO1_1_S151_<br>L006_R1_001_screen | PAO1 | 99967 | 75714 | 75.74 | 1286 | 1.29 |
| 6-PAO1_2_S152_<br>L006_R1_001_screen | PAO1 | 100023 | 98065 | 98.04 | 1809 | 1.81 |
| 7-CL PAO1_1_S153_<br>L006_R1_001_screen | CL PAO1 | 100007 | 53252 | 53.25 | 1654 | 1.65 |
| 8-CL PAO1_2_S154_<br>L006_R1_001_screen | CL PAO1 | 100075 | 58430 | 58.38 | 1707 | 1.71 |

**Table 4.**
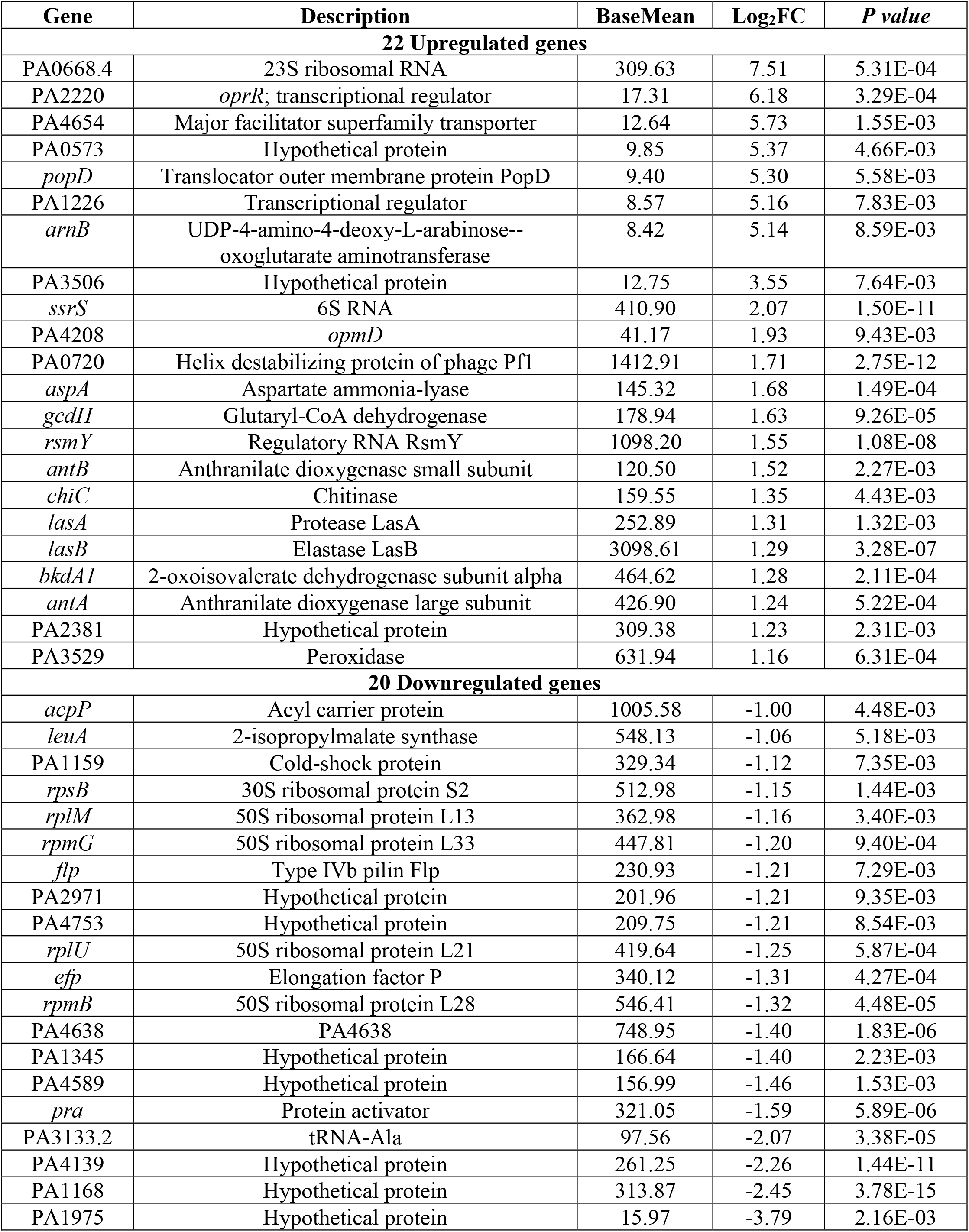
Differential expression of 42 *P. aeruginosa* genes on the cornea 4 h after bacterial challenge comparing prior lens wearing corneas (24 h) with naive corneas (also see Supplemental Table S7).

| Gene | Description | BaseMean | Log<sub>2</sub>FC | P value |
| --- | --- | --- | --- | --- |
| **22 Upregulated genes** |  |  |  |  |
| PA0668.4 | 23S ribosomal RNA | 309.63 | 7.51 | 5.31E-04 |
| PA2220 | *oprR*; transcriptional regulator | 17.31 | 6.18 | 3.29E-04 |
| PA4654 | Major facilitator superfamily transporter | 12.64 | 5.73 | 1.55E-03 |
| PA0573 | Hypothetical protein | 9.85 | 5.37 | 4.66E-03 |
| *popD* | Translocator outer membrane protein PopD | 9.40 | 5.30 | 5.58E-03 |
| PA1226 | Transcriptional regulator | 8.57 | 5.16 | 7.83E-03 |
| *arnB* | UDP-4-amino-4-deoxy-L-arabinose--oxoglutarate aminotransferase | 8.42 | 5.14 | 8.59E-03 |
| PA3506 | Hypothetical protein | 12.75 | 3.55 | 7.64E-03 |
| *ssrS* | 6S RNA | 410.90 | 2.07 | 1.50E-11 |
| PA4208 | *opmD* | 41.17 | 1.93 | 9.43E-03 |
| PA0720 | Helix destabilizing protein of phage Pf1 | 1412.91 | 1.71 | 2.75E-12 |
| *aspA* | Aspartate ammonia-lyase | 145.32 | 1.68 | 1.49E-04 |
| *gcdH* | Glutaryl-CoA dehydrogenase | 178.94 | 1.63 | 9.26E-05 |
| *rsmY* | Regulatory RNA RsmY | 1098.20 | 1.55 | 1.08E-08 |
| *antB* | Anthranilate dioxygenase small subunit | 120.50 | 1.52 | 2.27E-03 |
| *chiC* | Chitinase | 159.55 | 1.35 | 4.43E-03 |
| *lasA* | Protease LasA | 252.89 | 1.31 | 1.32E-03 |
| *lasB* | Elastase LasB | 3098.61 | 1.29 | 3.28E-07 |
| *bkdA1* | 2-oxoisovalerate dehydrogenase subunit alpha | 464.62 | 1.28 | 2.11E-04 |
| *antA* | Anthranilate dioxygenase large subunit | 426.90 | 1.24 | 5.22E-04 |
| PA2381 | Hypothetical protein | 309.38 | 1.23 | 2.31E-03 |
| PA3529 | Peroxidase | 631.94 | 1.16 | 6.31E-04 |
| **20 Downregulated genes** |  |  |  |  |
| *acpP* | Acyl carrier protein | 1005.58 | -1.00 | 4.48E-03 |
| *leuA* | 2-isopropylmalate synthase | 548.13 | -1.06 | 5.18E-03 |
| PA1159 | Cold-shock protein | 329.34 | -1.12 | 7.35E-03 |
| *rpsB* | 30S ribosomal protein S2 | 512.98 | -1.15 | 1.44E-03 |
| *rplM* | 50S ribosomal protein L13 | 362.98 | -1.16 | 3.40E-03 |
| *rpmG* | 50S ribosomal protein L33 | 447.81 | -1.20 | 9.40E-04 |
| *flp* | Type IVb pilin Flp | 230.93 | -1.21 | 7.29E-03 |
| PA2971 | Hypothetical protein | 201.96 | -1.21 | 9.35E-03 |
| PA4753 | Hypothetical protein | 209.75 | -1.21 | 8.54E-03 |
| *rplU* | 50S ribosomal protein L21 | 419.64 | -1.25 | 5.87E-04 |
| *efp* | Elongation factor P | 340.12 | -1.31 | 4.27E-04 |
| *rpmB* | 50S ribosomal protein L28 | 546.41 | -1.32 | 4.48E-05 |
| PA4638 | PA4638 | 748.95 | -1.40 | 1.83E-06 |
| PA1345 | Hypothetical protein | 166.64 | -1.40 | 2.23E-03 |
| PA4589 | Hypothetical protein | 156.99 | -1.46 | 1.53E-03 |
| *pra* | Protein activator | 321.05 | -1.59 | 5.89E-06 |
| PA3133.2 | tRNA-Ala | 97.56 | -2.07 | 3.38E-05 |
| PA4139 | Hypothetical protein | 261.25 | -2.26 | 1.44E-11 |
| PA1168 | Hypothetical protein | 313.87 | -2.45 | 3.78E-15 |
| PA1975 | Hypothetical protein | 15.97 | -3.79 | 2.16E-03 |

## Discussion

The aim of this study was to determine the impact of prior contact lens wear on the transcriptomic response of a healthy corneal epithelium to subsequent challenge with the opportunistic pathogen *P. aeruginosa*. Using a unique lens-wearing murine model that we have previously described (32), we compared gene expression in the corneal epithelium of mice that had previously worn a contact lens for 24 h with naïve (healthy) epithelium in their response to a 4 h challenge with *P. aeruginosa* (PAO1). Analysis of total RNA-sequencing revealed that some genes were altered in expression upon *P. aeruginosa* challenge only if the cornea had previously worn a lens (i.e. they were not changed by inoculation when there was no prior lens wear). This included upregulation of pattern-recognition receptor signaling genes (*tlr3, nod1*) and downregulation of the lectin pathway of complement activation (*masp1*). There were other genes impacted <u>differently</u> by inoculation if there was prior lens wear, including further upregulation of genes involved in innate (*gpr55*, *ifi205*, *wfdc2*) and acquired (*tcf7*) defense, and further downregulation of *efemp1* (required for corneal stromal integrity).

The study design also allowed us to determine the impact of lens wear alone on gene expression in the corneal epithelium, and how prior lens wear impacts gene expression in *P. aeruginosa* when it is exposed to the ocular surface. Lens wear upregulated innate and acquired immune defense genes (*axl, grn, tcf7, gpr55*) and downregulation of calcium-dependent genes (*necab1, snx31, npr3*) involved in cell signaling and cell sorting. Being exposed to a contact lens wearing cornea versus a naïve cornea caused a greater upregulation of *P. aeruginosa* genes involved in virulence (*popD*) and its regulation (*rsmY*, PA1226) and in genes involved in antimicrobial resistance (*arnB, oprR*).

Enrichment of pattern recognition receptor genes *tlr3* and *nod1* after post-lens wear challenge with *P. aeruginosa* suggests that lens wear may enhance the innate immune ‘tone’ of corneal epithelium responses to bacteria. TLR3 is expressed only in the epithelium of human cadaver corneas, and TLR3 agonists increase the expression of human cathelicidin (hCAP-18), an anti-*Pseudomonal* antimicrobial peptide, in primary cultured human corneal epithelial cells (75). Similarly, Nod1 (and Nod2) are well-recognized pattern recognition receptors that respond to microbial challenges (e.g. bacterial, viral) and sense danger signals associated with disruptions to cellular homeostasis (76,77). Thus, one effect of 24 h lens wear may be to ‘prime’ the epithelium to defend against microbial challenge which could be beneficial in preventing adhesion and subsequent infection. Conversely, binding PRRs could also activate local inflammation with the potential to promote epithelium susceptibility to infection.

Downregulation of *masp1* by *P. aeruginosa* challenge following 24 h lens wear versus naïve corneal epithelium suggests that lens wear could also compromise certain innate defenses of the cornea and ocular surface. *Masp1* encodes a mannose-binding lectin (MBL)-associated serine protease required to activate the lectin pathway of the complement system (47) and alternate pathway of complement *via* other MASPs (78). MBL is a pattern recognition molecule that recognizes carbohydrate residues, e.g. D-mannose, and other pathogen-associated molecular patterns, on microbial surfaces leading to activation of associated serine proteases (MASP1/MASP2) with C3 convertase (C4b2a) formation (79). Gene polymorphisms in the lectin pathway of complement are associated with early airway colonization by *P. aeruginosa* in cystic fibrosis (80). Since functionally-active complement proteins are present in human tear fluid, e.g. C3, C4 and Factor B (81), *masp1* down-regulation may hinder complement-mediated defense. However, given the role of complement in mediating pathology of some contact lens-associated complications (82), further study is needed to determine the significance of this finding.

Among the immune defense genes further upregulated in response to *P. aeruginosa* after a lens had been worn, the largest difference was in *tcf7*, encoding a T cell-specific transcription factor Tcf1, along with three other notable genes associated with (or predicted to be involved with) innate immunity (*gpr55, wfcd2, ifi205*). Tcf1 has multiple roles in T cell function in health and disease, e.g. T cell development, memory cell formation, immune regulation (83). Greater upregulation of *tcf7* in response to bacteria after lens wear may be relate to the finding that T cells are present in human corneas and interact with sensory nerves and other resident immune cells (84). However, with multiple known and potential functions of Tcf1 (positive and negative regulation of immune responses), it is difficult at present to predict the impact of this change. Nevertheless, along with enhanced expression of *gpr55* involved in neutrophil recruitment after injury (51) and of *ifi205* regulated by the type 1 interferon (IFN-β), these changes are consistent with 24 h of lens wear ‘priming’ the corneal epithelium for enhanced immune defense against infection.

Among the genes further downregulated by *P. aeruginosa* after lens wear was *efemp1*, encoding epidermal growth factor (EGF)-containing fibulin-like extracellular matrix protein 1 (also known as Fibulin-3), found in basement membranes of many tissues. *Efemp1* is expressed in the cornea and other structures in the eye, and gene-knockout mice loss (*efemp1*^-/-^) show corneal dysfunction with stromal thinning at 2 months of age and corneal opacity and vascularization at 9 months (55). In other tissues, EFEMP1 inhibits the growth of carcinoma cells and promotes their apoptosis (54). Suppression of *efemp1* in response to *P. aeruginosa* by prior lens wear could be significant in the development of corneal infections given the role of the basement membrane in epithelial resistance to *P. aeruginosa* traversal (85). The impact of *P. aeruginosa* alone compared to naïve corneas also revealed interesting changes in gene expression in the corneal epithelium. They included upregulation of *klf10* and *fxyd1* which regulate cell differentiation and protect vasculature against oxidative stress respectively (56,57,86) and downregulation of *kat6A* a lysine acetyl-transferase (58) and *jag1* a ligand for Notch signaling required for normal ocular lens development (59). Further network analysis revealed an unexpected over- representation of mitochondrial genes identified in response to *P. aeruginosa* that may suggest involvement in maintaining tissue homeostasis and resistance to bacterial exposure. Other changes in epithelium response to *P. aeruginosa* alone grouped into categories of neurotrophin binding and protein stabilization. Each of these changes are likely to be of interest in the context of corneal homeostasis and intrinsic defense against bacteria and would be worthy of further follow-up in future studies.

ClueGO analysis of the impact of lens wear alone on corneal epithelium clustered some of the most upregulated genes (*axl*, *gpr55*, *tcf7*, *klf10*) into a network of “regulation of leukocyte differentiation”. For example, *axl* is a negative regulator of TLR-mediated signaling and associated cytokine expression (e.g. involving dendritic cells) serving to limit inflammation after pathogen or antigen recognition (87). Conversely, *gpr55* (further upregulated by *P. aeruginosa* challenge, see above) encodes a G-protein coupled receptor that binds proinflammatory lipids (lysophosphatidylinositols) (88) is not only associated with neutrophil recruitment (51) but other proinflammatory effects *via* expression on monocytes and Natural Killer cells (89). Upregulation of these genes by lens wear correlates with our observed induction of parainflammation in the cornea after 24 h wear, the latter involving CD11c+, Lyz2+ and MHC-II+ cells (32,43,90,91) some of which are likely to be dendritic cells and/or monocytes. Longer duration of lens wear also recruits Ly6G+ cells (likely neutrophils) via ψο T cells and IL-17A (92). Thus, an examination of the role of *axl* and *gpr55* in lens-induced corneal parainflammation would be of value in future studies. Some known roles of *tcf7* and *klf10* were mentioned above in the context of epithelial responses to *P. aeruginosa*. Given that both were also upregulated by lens wear alone, it is possible that both also influence corneal parainflammation during and after lens wear. Indeed, *klf10* is induced by TGF-ß to suppress inflammation, slow host cell proliferation and induce cellular apoptosis. Interestingly, contact lens wear in rabbits also reduces corneal epithelial cell proliferation albeit with RGP lenses (37).

*Grn*, encoding progranulin, was identified in the corneal epithelium using single-cell RNA sequencing (93). Lens wear alone significantly upregulated *grn* expression in our study. While the function of *grn* has not been studied in the cornea, progranulin knockout mice (*grn*-/-) show greater infiltration of *iba1+* cells (CD68+), increased expression of proinflammatory molecules (TNF-𝛼, IL-1ß, C3, CCL2) and increased VEGF-A in a macrophage cell line under hypoxic conditions (94). As such, the role of *grn* in lens-induced parainflammation would seem a worthwhile avenue of future study.

Lens wear alone also downregulated a number of calcium-dependent genes (*necab1, snx31, npr3*). None of these genes have been studied in the context of corneal physiology, however their functions in other organ systems suggest that they may play important roles in the cornea to regulate tissue homeostasis and inflammation. For example, *snx31* encodes a novel sorting nexin associated with endocytic trafficking and potential degradation of apical surface uroplakins in bladder urothelium (95), and its association with terminally-differentiated cells suggests involvement in regulation of urothelial barrier function. Thus, *snx31* downregulation by lens wear alone represents a potential avenue of investigation of lens-mediated disruption of corneal homeostasis.

The impact of prior lens wear on *P. aeruginosa* gene expression could also be significant to our understanding of the pathogenesis of infectious keratitis and other lens wear associated complications. For example, prior lens wear upregulated *P. aeruginosa popD*, a T3SS translocon protein. The T3SS is well established as a major contributor to *P. aeruginosa* virulence in the cornea (68,69,96,97), with *popD* alone modulating host cell function (64). Similarly, *P. aeruginosa* survival at the corneal surface could be promoted by upregulation of *arnB*, part of the *arn* operon that facilitates resistance to polymyxin B and host defense antimicrobial peptides, e.g. β-defensins, which share similar properties and mechanism of action (15). Indeed, we showed human tear fluid down-regulates the *arn* operon correlating with increased susceptibility to polymyxin B (29) suggesting a role in tear-mediated defense that may be compromised by prior lens wear. Upregulation of PA1226 after encoding *oprR* which mediates resistance to quaternary ammonium compounds (74) and contributes to *P. aeruginosa* biofilm formation and *in vivo* virulence (98), will also likely to favor bacterial survival as the ocular surface and in the face of contact lens care solutions. Another change that favors *P. aeruginosa* survival and adaptation was upregulation of *rsmY* encoding a small regulatory (non-coding) RNA (61,62). Together, these data suggest that prior lens wear can alter how *P. aeruginosa* responds to the cornea moving towards phenotypes that can favor survival, adaption, persistence and virulence.

While the current study revealed a large amount of information contributing to our limited understanding of lens wear impacts, further work will be needed to follow up this study. This includes analysis of corneal epithelium protein expression and functional changes to epithelial homeostasis after lens wear with and without *P. aeruginosa* challenge to determine their significance to the pathogenesis of infection and other lens-associated adverse events. It would also be of interest to perform single cell RNA- sequencing to help determine relative contributions of different cell types within the corneal epithelium, e.g. resident immune cells and different epithelial cell types, along with the influence of other cells impacting corneal homeostasis, e.g. stromal keratocytes and sensory nerves. This study evaluated impact of a single time point (24 h) of lens wear, followed by lens removal, the latter done to minimize confounding issues likely to arise if a lens remains in place during bacterial challenge, e.g. effects a lens in place on bacterial adherence and corneal responses to them. Here, we deliberately removed those confounders so we could tease out the contribution of one aspect of what is likely to be a complex interplay between the cornea, bacteria, the lens and other factors present *in vivo*. Further studies could examine bacterial-epithelium interactions with a lens in place and other time points.

In summary, this study provides insights into the impact of contact lens wear on the corneal response to *P. aeruginosa,* to the lens itself, and on bacterial gene expression when they are inoculated onto a cornea. Using corneal health as the baseline and future destiny in at least three of the four experimental groups (naïve, lens alone, inoculated alone) highlighted transcriptional responses in the epithelium that help restore corneal homeostasis or trigger an alternate homeostasis that predisposes to infection or other lens-associated adverse events. The outcomes illustrate the complexity of impacts of a medical device on the host, the pathogen, and potentially on host-pathogen interactions. Impacted genes and associated networks identified provide targets for future study and for hypothesis development towards a better understanding of corneal responses to lens wear that can result in infectious or other adverse pathology.

## Acknowledgements

This work was supported by the National Eye Institute EY030350 (SMJF). Thanks to Dr. Abby Kroken, now at Loyola University, Chicago IL, for contributing to the artwork schematic in Figure 1.

---

Figure S1. Transcriptional changes in the naïve corneal epithelium after 4 h exposure to *P. aeruginosa*. A) Enrichment map categories of differentially-expressed genes. Node size is proportional to enrichment score (NES= 0.5 – 1). B) Network map of genes in the ubiquinone network that were enriched after bacterial challenge. Nodes are annotated with the Log_2_ Fold-Change. Blue = upregulated, Red = downregulated.

Figure S2. Gene ontology enrichment map of differentially-expressed genes in a naïve corneal epithelium after 4 h exposure to *P. aeruginosa* that were not captured by an immune pathway analysis. Nodes are annotated with the Log_2_ Fold-Change. Blue = upregulated, Red = downregulated.

**Supplemental Table S1.** Deregulated genes (224) in the mouse corneal epithelium unique to 24 h lens wear then 4 h exposure to *P. aeruginosa* versus naïve control (no lens wear, no inoculation).

**Supplemental Table S2.** (A) Mouse corneal epithelium genes (143) deregulated in response to 4 h *P. aeruginosa* exposure in both prior lens wearing and naïve corneas. (B) Relative fold-change of the most deregulated genes comparing response to *P. aeruginosa* with and without prior lens wear.

**Supplemental Table S3.** Deregulated genes (131) in the mouse corneal epithelium unique to 4 h exposure to *P. aeruginosa* versus a naïve cornea.

**Supplemental Table S4.** Deregulated genes (94) in the murine corneal epithelium after 24 h lens wear versus a naïve cornea. See network analysis in Fig. 4A

**Supplemental Table S5.** Sets of deregulated genes involved in the murine corneal epithelium response to 24 h of lens wear alone. See enrichment map in Fig. 4B.

**Supplemental Table S6.** ClueGo analysis of immune response-related genes involved in murine corneal epithelium response to 24 h lens wear alone. See Fig. 4C.

**Supplemental Table S7.** Deregulated bacterial genes (183) after 4 h exposure to the murine corneal epithelium comparing prior lens wear (24 h) to a naïve control. See Table 4.

