## Supplementary figures and images for "Contact Lens Wear Alters Transcriptional Responses to *Pseudomonas aeruginosa* in Both the Corneal Epithelium and the Bacteria"

### Supplemental Fig. S1

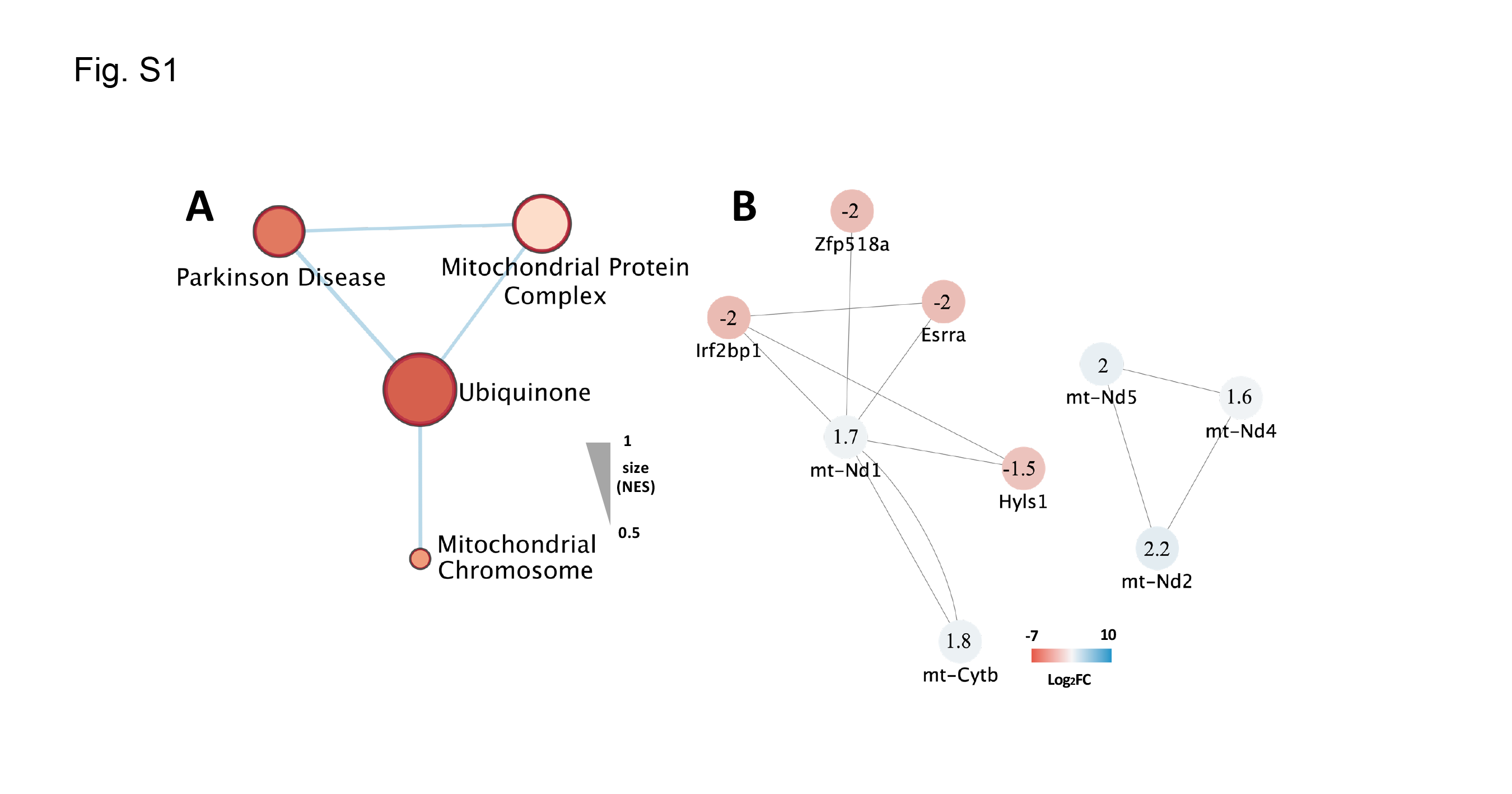

### Supplemental Fig. S2

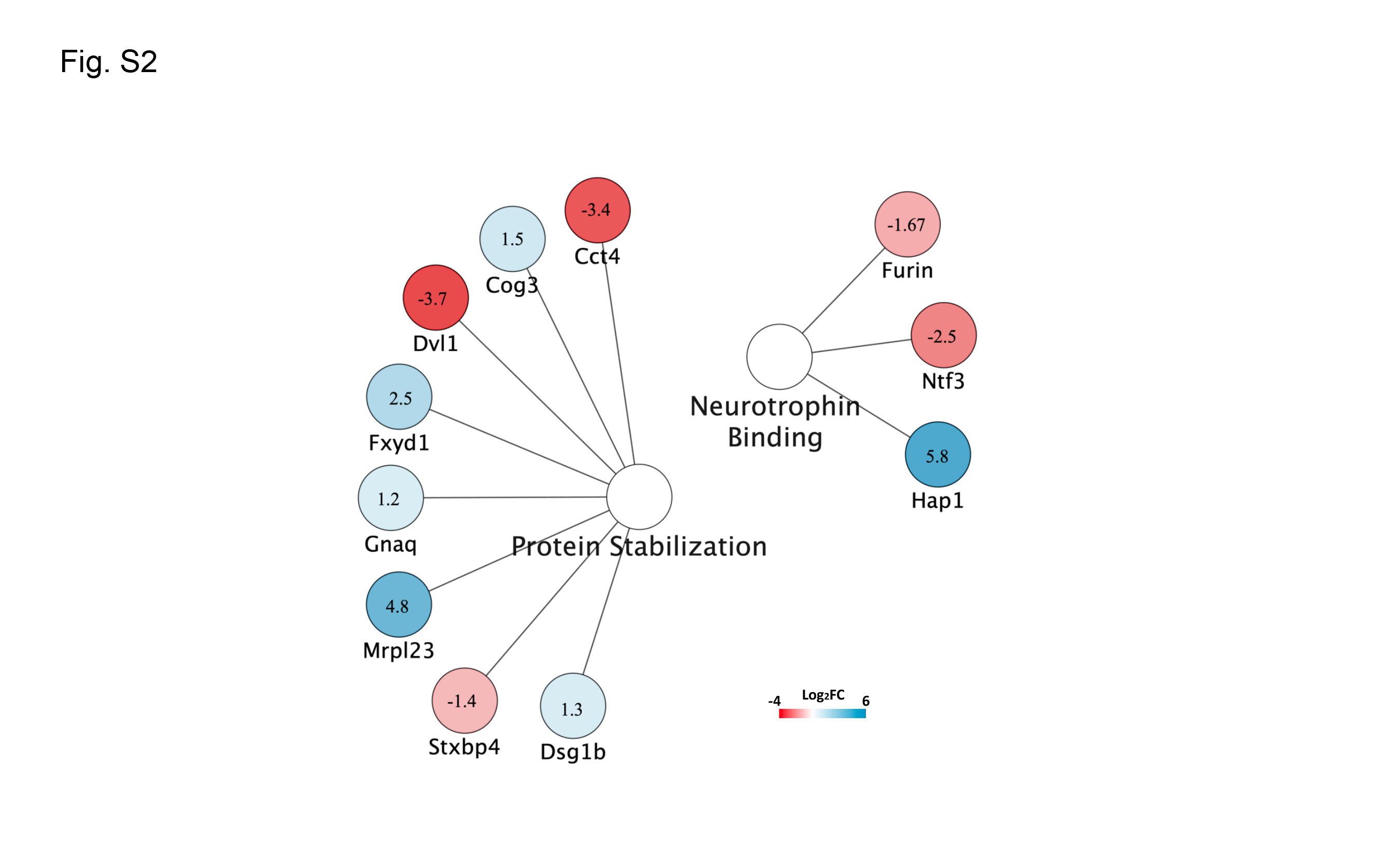
